# H4K20me1 plays a dual role in transcriptional regulation during regeneration and axis patterning of *Hydra*

**DOI:** 10.1101/2022.06.01.494139

**Authors:** Akhila Gungi, Mrinmoy Pal, Shagnik Saha, Sanjeev Galande

## Abstract

The evolution of the first body axis in the animal kingdom and an extensive ability to regenerate makes *Hydra*, a Cnidarian, an excellent model system for understanding the underlying epigenetic mechanisms. We identify that SETD8 is critical for regeneration due to its interaction with β-catenin to fine-tune the underlying gene regulatory network. Its target histone mark, H4K20me1, colocalizes with transcriptional activation machinery locally at the β-catenin bound TCF/LEF binding sites on the promoters of head-associated genes, marking an epigenetic activation node. Contrastingly, genome-wide analysis of the H4K20me1 occupancy revealed a negative correlation with transcriptional activation. We propose H4K20me1 as a general repressive histone mark in Cnidaria and describe its dichotomous role in transcriptional regulation in *Hydra*.

## INTRODUCTION

Post-translational modifications (PTMs) of proteins are major regulators of cellular functions (Deribe et al. 2010), with nuclear histone PTMs being prominent context-specific players in transcriptional regulation. Although the methylation of histone H4 was one of the first histone PTM discovered (DeLange et al. 1969), the modifiers responsible are recent discoveries. While several enzymes are known to deposit the di- and tri-methyl marks on H4K20 — SUV4-20H1 and SUV4-20H2 being the predominant ones (Schotta et al. 2008), there is only one known monomethyltransferase — SETD8 (KMT5A or Pr-SET7) (Oda et al. 2009). SETD8 has many functions in cells and acts by disrupting signaling pathways (Ke et al. 2014), regulating transcription factors (Choi et al. 2020), altering the chromatin accessibility around the promoters of genes (Myers et al. 2020), and preventing both oncogene-induced and replicative cellular senescence by suppressing nucleolar and mitochondrial activity (Tanaka et al. 2017). SETD8 is involved directly in the Wnt/β-catenin signaling pathway in mammalian cells, *Drosophila* larvae, and zebrafish (Li et al. 2011). SETD8 is required to activate Wnt target genes by interaction with TCF4 during the development of zebrafish and the wings of *Drosophila* (Li et al. 2011; Yu et al. 2019). The methylation of H4 is highly evolutionarily conserved and exists in three states as mono-, di-, and trimethylation. While the mono- (H4K20me1) and di-methylated (H4K20me2) H4K20 are involved in DNA replication and DNA damage repair, the trimethylated H4K20 (H4K20me3) is a mark of silenced heterochromatic regions (Schotta et al. 2004; Schotta et al. 2008). The function of H4K20me1 is elusive, with reports of activation (Kapoor-Vazirani and Vertino 2014; Lv et al. 2016; Shoaib et al. 2021) and repression (Nishioka et al. 2002; Tjalsma et al. 2021) of transcription.

To understand the role of H4K20me1 in transcriptional regulation of developmental processes, including axis patterning and regeneration, *Hydra* serves as an excellent model organism. *Hydra* belongs to the phylum Cnidaria which is the phylum that innovated a body axis during the evolution of multicellular organisms. It also harbours extensive powers of regeneration which require the recapitulation of the axis patterning gene regulatory networks to be formed *de novo*. Both embryonic and regenerative *de novo* axis patterning is regulated by the head organizer Wnt/β-catenin signaling pathway. While multiple studies have investigated the molecular networks underlying the axis patterning (Bode 2011; Gufler et al. 2018; Reddy et al. 2019; Vogg et al. 2019; Moneer et al. 2021; Unni et al. 2021), few study the epigenetic regulators of gene expression (Reddy et al. 2020). We identify that the H4K20me1 writer SETD8 is critical for Wnt-triggered regeneration and that the Wnt pathway regulates both the enzyme and the target modification. We also identify dual modes of transcriptional regulation by H4K20me1, activating transcription at promoters downstream to specific signaling pathways. Additionally, at the whole genome level, it excludes activation-associated features allowing us to describe the presence of a repressive histone mark for the first time in Cnidaria.

## RESULTS

### SETD8 is important for both head and foot regeneration in Hydra

In *Hydra*, upon amputation, the injury site triggers a wound healing response involving re-organization of the epithelial cells in 1 hour. Following this, morphological changes are only visible from 30 hours post-amputation (hpa), and at 30–36 hpa, tentacle buds start emerging, indicating successful differentiation of head structures. The emergence completes over the next 24 h and by 72 hpa, fully functional tentacles and hypostome are formed. Basal disk upon amputation regenerates within 30–36 hpa. The morphological characteristics of a head regenerating polyp are depicted in Fig. 1A. Upon a screen using specific pharmacological inhibitors (Fig. 1B, Fig. S1), we identified a significant role for SETD8 in head and foot regeneration. When KMT5A (SETD8) was attenuated, a significant reduction in the head regenerative ability in *Hydra* polyps was observed at all the target timepoints starting from 33 hpa. The emergence of tentacles is delayed by 12 h, and the polyps fail to successfully regenerate all their tentacles at the same time as control polyps (Fig. 1C, 1D). Since SETD8 inhibition significantly impacts head regeneration, we performed a foot regeneration assay to determine the role of SETD8 in this process. Following amputation of the foot, a peroxidase staining assay was employed to understand the dynamics of the regenerative process. As seen in Fig. 1E, the control polyps gradually form a differentiated foot following amputation, starting from 26 hpa and completed by 36 hpa. Contrastingly, the inhibitor-treated polyps cannot regenerate the foot after amputation (Fig. 1E).

**Fig. 1:**
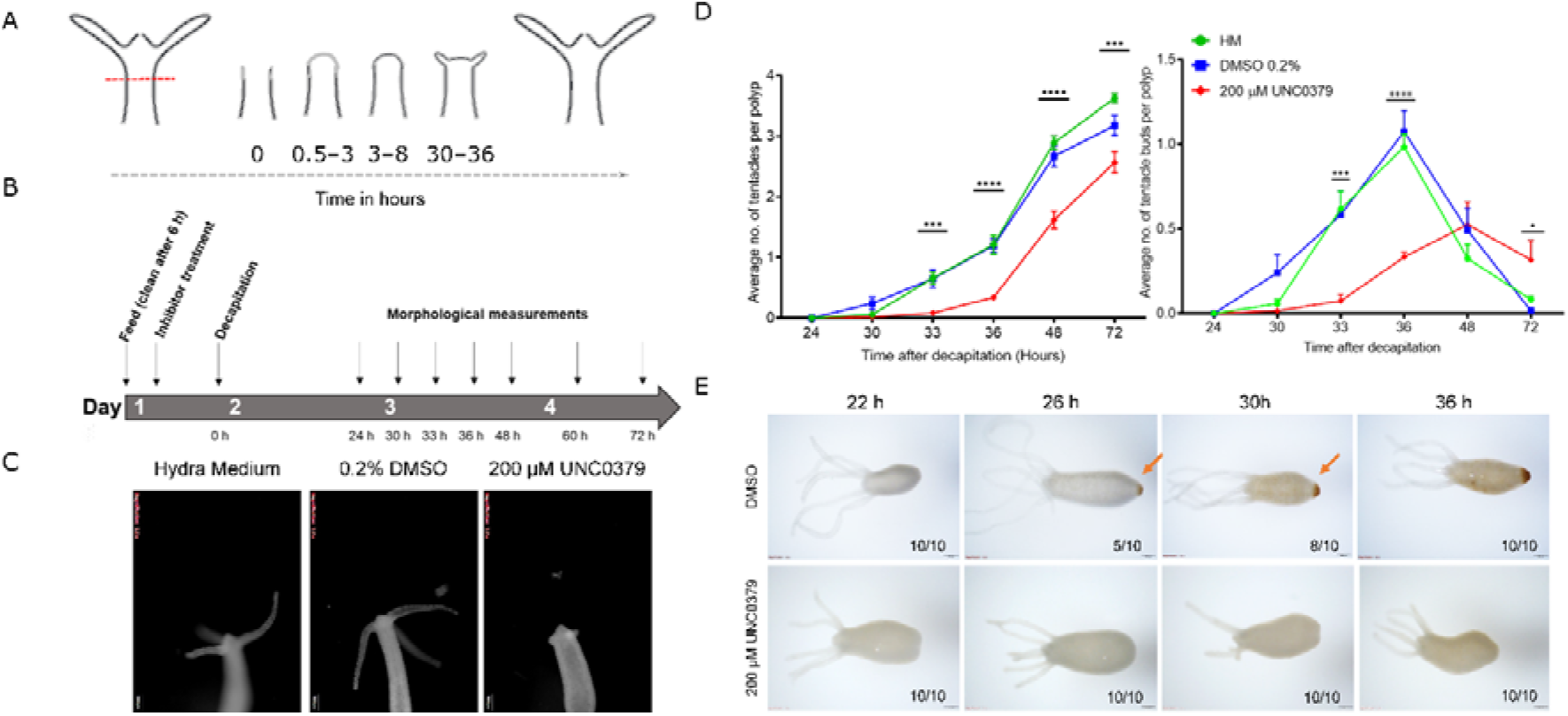
SETD8 activity is necessary for regeneration in *Hydra*. A) Kinetics of gross morphological changes during *Hydra* head regeneration. B) Regime for the chemical inhibitor treatment and head regeneration assay. C) Polyps decapitated and fixed at 72 hours post-amputation (hpa) with and without treatment with the SETD8 inhibitor, UNC0379. D) The graph depicts the average number of tentacles (left) and the number of tentacle buds per polyp (right) at each time point post decapitation during regeneration. (N = 5, n = 25, *- P <0.5, ***- P < 0.001, ****- P <0.0001). E) A regeneration time course for observing the process of foot regeneration following amputation. Polyps fixed at 22 h, 26 h, 30 h, and 36 hpa were subjected to peroxidase-staining assay using DAB, resulting in a brown precipitate at the regenerated foot. Upon treatment with the inhibitor UNC0379, foot regeneration is severely impaired. The scale bar measures 200 μm.

### SETD8 regulates head organizer gene expression during regeneration

The wound healing process is the first step of regeneration, and its dynamics were studied following the inhibition of SETD8. We performed staining of the F-actin filaments in the regenerating tip using phalloidin and observed the cytoskeletal structures at early time points post decapitation. There was no significant difference in the extent of wound closure at comparable time points between the control and inhibitor-treated polyps (Fig. 2A). Following successful wound healing, the head organizer is formed in *Hydra*, which is needed for the differentiation of the head structures like mouth and tentacles. In *Hydra*, the Wnt signalling pathway and activity of β-catenin are required in the early stages of regeneration of both the head and the foot (Gufler et al. 2018). To understand further molecular dynamics, we performed 3’mRNA sequencing on the regenerating tips of polyps treated with UNC0379 after different regeneration times. We identified various signalling pathway members, among which the Wnt signalling pathway components were enriched (Fig. 2B). Among the various genes that are part of the Wnt signalling pathway and are direct targets of the TCF7L2 transcription factor, *Brachyury* has been established as a bonafide head specification marker. *Brachyury* in *Hydra* is a head organizer gene involved in head morphogenesis. Therefore, we used this gene as a marker for the early morphogenetic events that take place post decapitation during regeneration. We performed a whole-mount RNA *in situ* hybridization to study the localization of the *Hv_Brachyury1* gene in the regenerating tips. In normal polyps, the expression of *Hv_Brachyury1* starts at 2 hpd at the regenerating tip. At 4 hpd, a scattered group of cells in the top 1/3^rd^ of the regenerating polyp start expressing this gene which later gets clustered to the tip of the animal by 8 hpd. This concentrated expression increases in intensity by 12 hpd and is maintained throughout following successful regeneration, as is seen in uncut normal polyps. When the polyps are subjected to the SETD8 inhibitor before decapitation, the expression of *Hv_Brachyury1* is severely affected, with both the expression domains and intensity reduced at all time points of regeneration (Fig. 2C). To understand the localisation of SETD8 and its target histone modification in *Hydra* polyps, we performed an immunofluorescence assay using specific antibodies against the methyltransferase and the histone PTM. The occurrence of both SETD8 and its target histone modification, H4K20me1, is relatively higher in the head and the foot of the polyps (Fig. 2D).

**Fig. 2:**
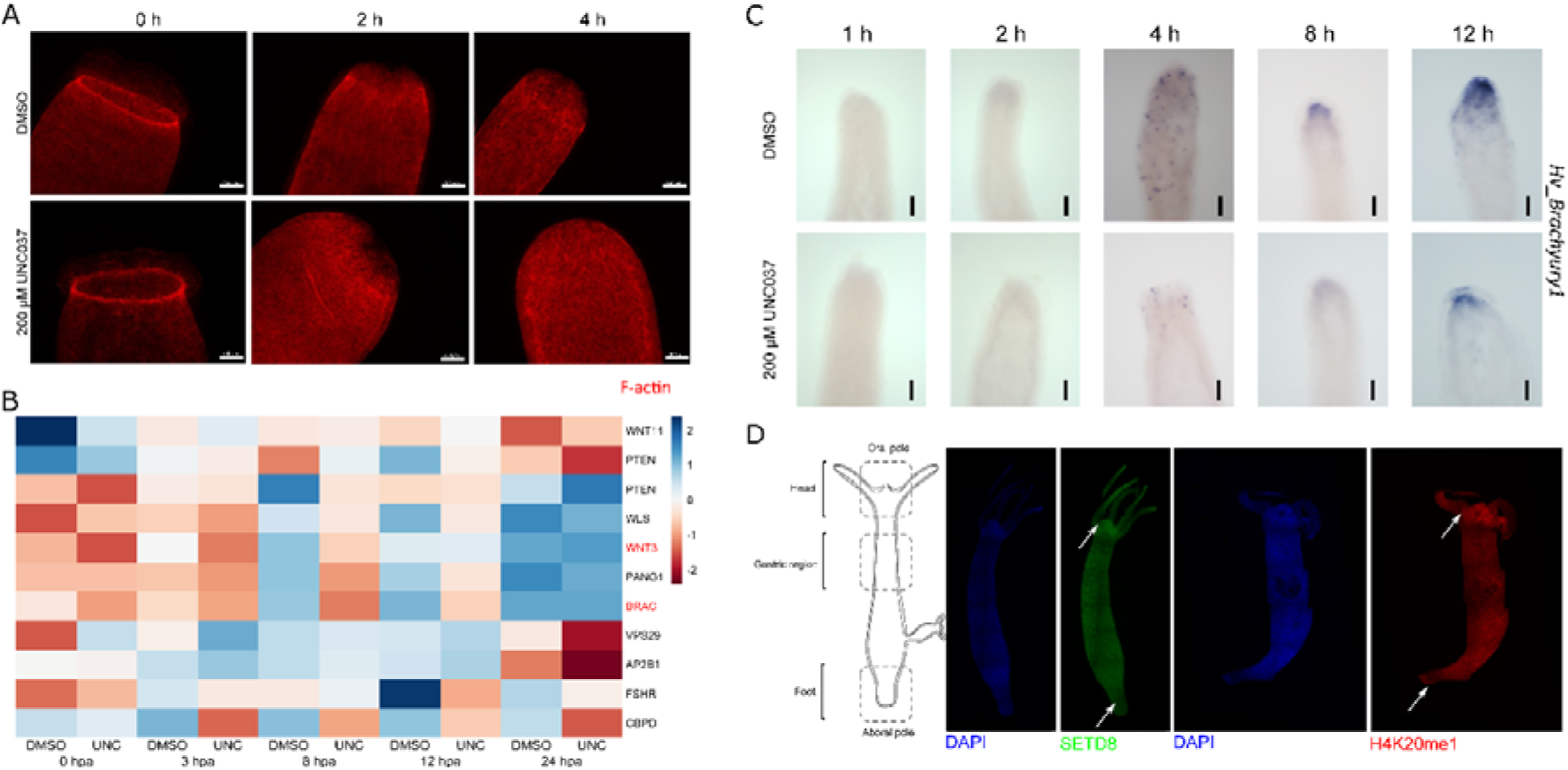
SETD8 regulates head organizer gene expression during regeneration. A) Wound healing during regeneration. The regenerating polyps at early time points are stained with phalloidin to observe the filamentous actin structures. The top row depicts control polyps, and the bottom row depicts the polyps treated with SETD8 inhibitor. The scale bar measures 200 μm. B) Differential expression of Wnt signaling-associated genes during head regeneration upon inhibition of SETD8. The plot depicts the expression in control and treated regenerating tips at each time point. Genes with a log2 fold change cut-off of ±0.58 and a P value of 0.01 have been depicted. C) Whole-mount RNA in situ hybridization against the Brachyury1 gene in regenerating polyps with and without inhibitor treatment. The blue stain depicts the expression pattern of the Brachyury1 gene and shows that the inhibitor treatment impairs the expression of Brachyury1. The scale bar measures 100 μm. D) The schematic of the *Hydra* polyp on the left depicts the three different regions of the polyp. The right panels depict polyps stained with α-SETD8 and α-H4K20me1 antibodies in red and green, respectively. The nuclei have been stained with DAPI as a counterstain.

### SETD8 interacts with the effector transcriptional activator of the Wnt signalling pathway in Hydra

*Brachyury1* is a target of the canonical Wnt/β-catenin signalling pathway in *Hydra*. To understand the role SETD8 has in the patterning of the oro-aboral axis of *Hydra*, we activated the Wnt signalling pathway using Alsterpaullone (ALP). Following the ALP treatment regime shown in Fig. 3A, which leads to the whole polyp turning into a head (Fig. 3B), we checked the levels of SETD8 at both the mRNA and protein levels. We observed that the expression of SETD8 is upregulated following the activation of the Wnt signalling pathway (Fig. 3C). The predicted size of *Hydra* SETD8 protein is 32.2 KDa, and as seen in Fig. 3D, the Western blot performed with an α-mouse-SETD8 antibody detects the target protein in the *Hydra* lysate. To further characterise the role of SETD8 in the Wnt signalling pathway, we performed a co-immunoprecipitation using an α-β-catenin antibody. Upon probing with a α-SETD8 antibody, we observed a clear pulldown of the SETD8 protein, indicating that SETD8 interacts with β-catenin in *Hydra* (Fig. 3E). Since SETD8 interacts with and is regulated by components of the Wnt/β-catenin regulatory network, we were interested in identifying the regulation of the target histone modification, H4K20me1. In a previous study, we identified a direct target of β-catenin named *Margin*. This is a homeodomain-containing transcription factor, which is upregulated upon activation of the Wnt signalling pathway and downregulated upon knockdown of β-catenin. β-catenin also has binding motifs at the promoter of this gene and binds upon its nuclearization when the signalling pathway is activated (Reddy et al. 2019). We used this experimental paradigm of ectopic activation of the Wnt signalling pathway and checked the occupancy of H4K20me1 on the promoter regions of *Margin* and *setd8*. We observed an enhanced occupancy of the modification at the promoter regions of both the genes (Fig. 3F).

**Fig. 3:**
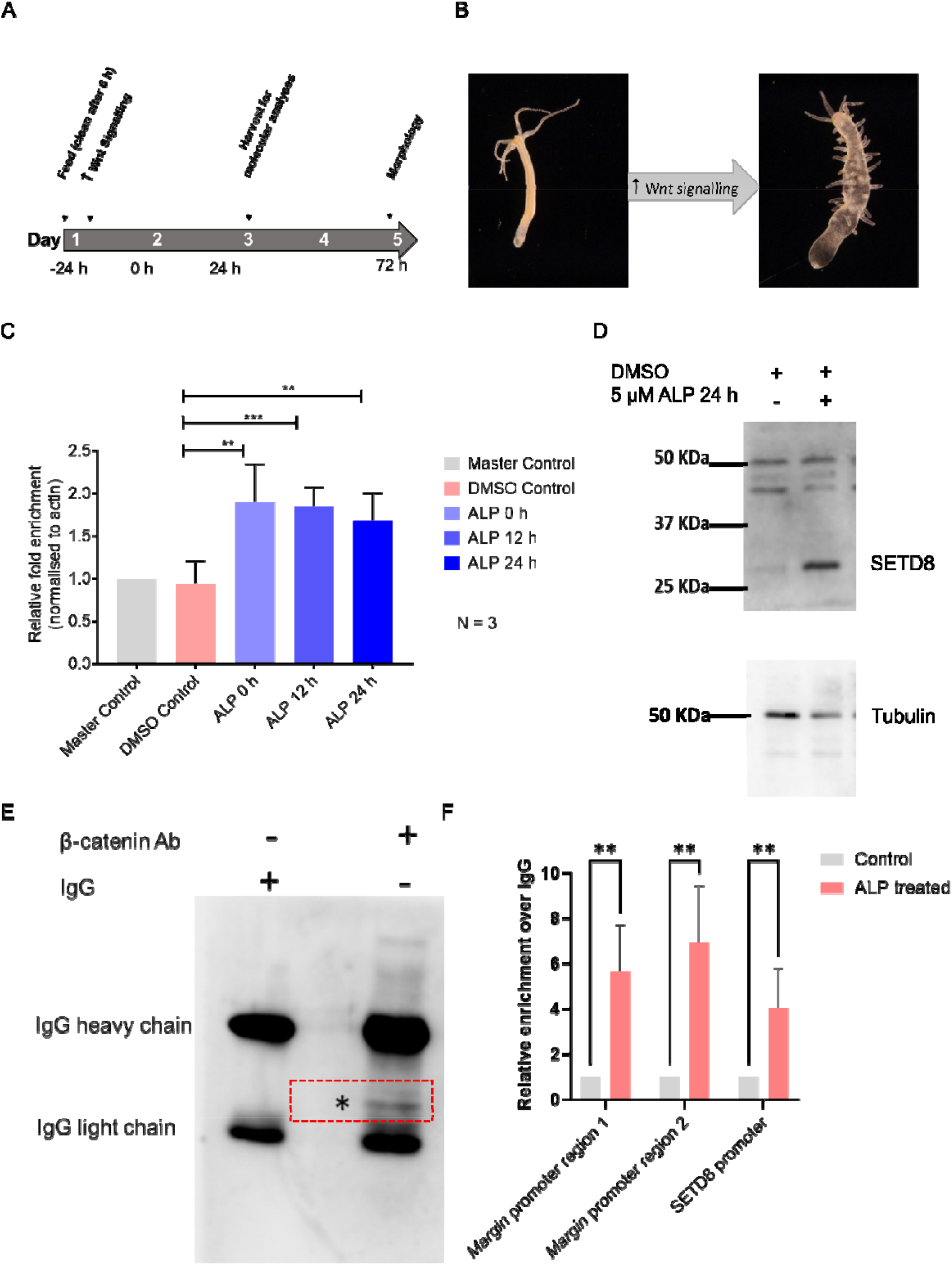
SETD8 is regulated by the Wnt/β-catenin signaling pathway and is part of the transcriptional machinery with β-catenin. A) The treatment regime used to activate the Wnt signaling pathway using ALP and study the level of SETD8. B) Following activation of Wnt signaling in *Hydra* the entire polyp turns into a head, and ectopic tentacles are seen all over the body column. C) The levels of SETD8 mRNA were investigated using qRT-PCR and show that the expression of SETD8 is upregulated following ALP treatment. D) The level of SETD8 protein was studied using Western blot, and this also shows the increased level upon activation of the Wnt signaling pathway. The size of *Hydra* SETD8 is 32.2 KDa, and the molecular weight ladder is indicated on the left of the gel. E) Western blot showing the co-immunoprecipitation of SETD8 upon pulldown using an α-active βcatenin antibody. The red box depicts the band corresponding to *Hydra* SETD8 protein which is 32.2 KDa in size, and the asterisks mark the SETD8 band in the lysates following immunoprecipitation. F) The occupancy of H4K20me1 was assayed using ChIP followed by q-PCR and plotted. The occupancy in the ALP-treated samples has been normalized to that in the DMSO control samples. (**- P = 0.0073, Two-way ANOVA with Šídák’s multiple comparisons test).

### H4K20me1 occupancy is context-dependent and is linked to specific signalling pathways

To elucidate the role of this unique modification in the regeneration of *Hydra*, we performed ChIP-sequencing for H4K20me1 in Wnt-activated polyps (Fig. 4) and regenerating tips at five time points (Fig. ***5***). The occupancy of H4K20me1 was plotted on gene bodies which are regions comprised of the coding sequence (CDS) and 2 Kb of genomic DNA on either ends of the CDS (Fig. 4A, Fig. 4B, Fig. ***5***A). K-means clustering revealed four clusters of gene bodies in the *Hydra* genome based on the occupancy of H4K20me1 in control and Wnt-activated polyps. The clusters 1 to 4 in H4K20me1 occupied regions of the ALP treated polyps contain 9240, 9579, 9245, and 23046 transcripts, respectively, including all the isoforms identified following in the latest version of the *Hydra* genome assembly. Based on the occupancy of H4K20me1 across the timepoints of regeneration, four k-means clusters were generated, which are shown in Fig. ***5***A. Clusters 1 to 4 contain 7514, 13251, 9867, and 20658 transcripts, respectively, including all the isoforms identified following in the latest version of the *Hydra* genome assembly. In both physiological conditions, the first cluster has a uniform distribution of the modification throughout the CDS, and this is the cluster with the maximum level of the modification too. The second and third clusters display occupancy of the mark on the 5’ end and the 3’ end of the gene body, respectively, with a distinct depletion on the other end. The fourth cluster has little to no occupancy of the H4K20me1 mark (Fig. 4B, Fig. ***5***A). Using the gene ontology (GO) terms associated with critical physiological processes, we plotted the relative enrichment of genes in the four clusters. While many signalling pathway genes displayed an occupancy of the H4K20me1 mark around the gene bodies, the Hippo signalling pathway had the least number of gene bodies with this modification (Fig. 4C, Fig. ***5***B). Since the role of H4K20me1 in transcriptional regulation is ambiguous or context-dependent, to understand the same, we used the publicly available datasets on regenerating *Hydra* and correlated our data with that. The cluster 1 genes with the highest level of H4K20me1 on the CDS, have the least occupancy of H3K4me2, H3K4me3, and H3K27ac at both the promoter regions and on the CDS. Cluster 2 genes with a low level of H4K20me1 at the promoter with a steep decline across the CDS, show a depletion of H3K4me2 on the CDS, and extremely low levels of the remaining two histone marks being investigated. The cluster 3 genes which have clear depletion of H4K20me1 at the promoter regions, show an enrichment of the activation-associated histone marks. The cluster 4 genes have the least occupancy of the H4K20me1 mark but show significant enrichment of the other three marks at the promoter regions (Fig. 4D, Fig. ***5***C).

**Fig. 4:**
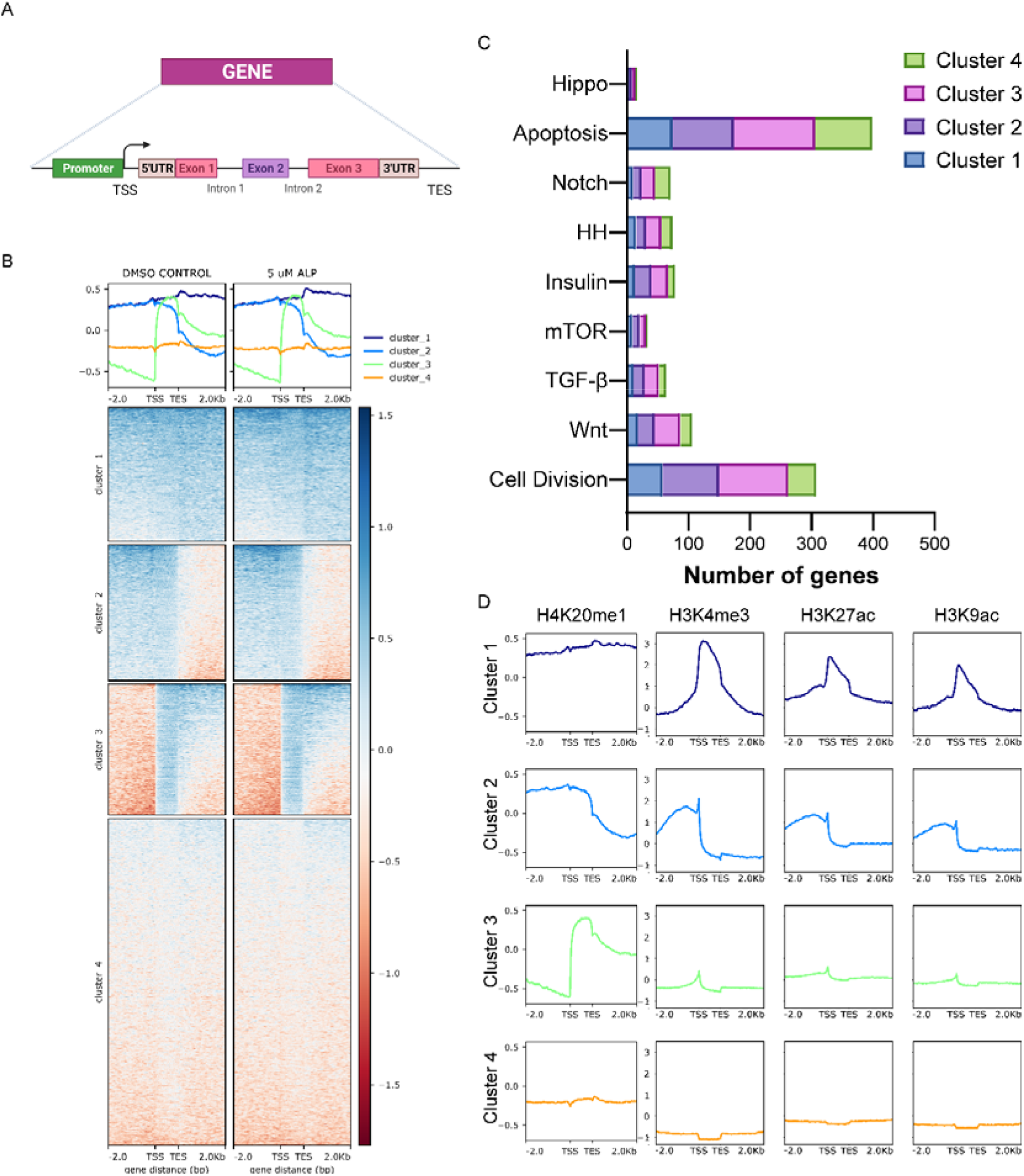
H4K20me1 occupancy is modestly increased globally on the gene bodies in *Hydra* upon activation of the Wnt/β-catenin signaling pathway. A) A schematic of the gene body that is used for H4K20me1 occupancy analyses. B) H4K20me1 ChIP signal on gene bodies of DMSO control and ALP treated (24h) polyps separated into 4 clusters. The occupancy of H4K20me1 on each gene body is shown in the clustered heatmaps (bottom panel). The profile plots (top panel) show the average ChIP signals of each cluster. C) Representation of pathways in the four clusters obtained based on the distribution of H4K20me1. D) The four clusters obtained based on the occupancy of H4K20me1 on gene bodies have been used to understand the occupancy of the three activation-associated histone marks: H3K4me3, H3K27ac, and H3K9ac in control polyps.

**Fig. 5:**
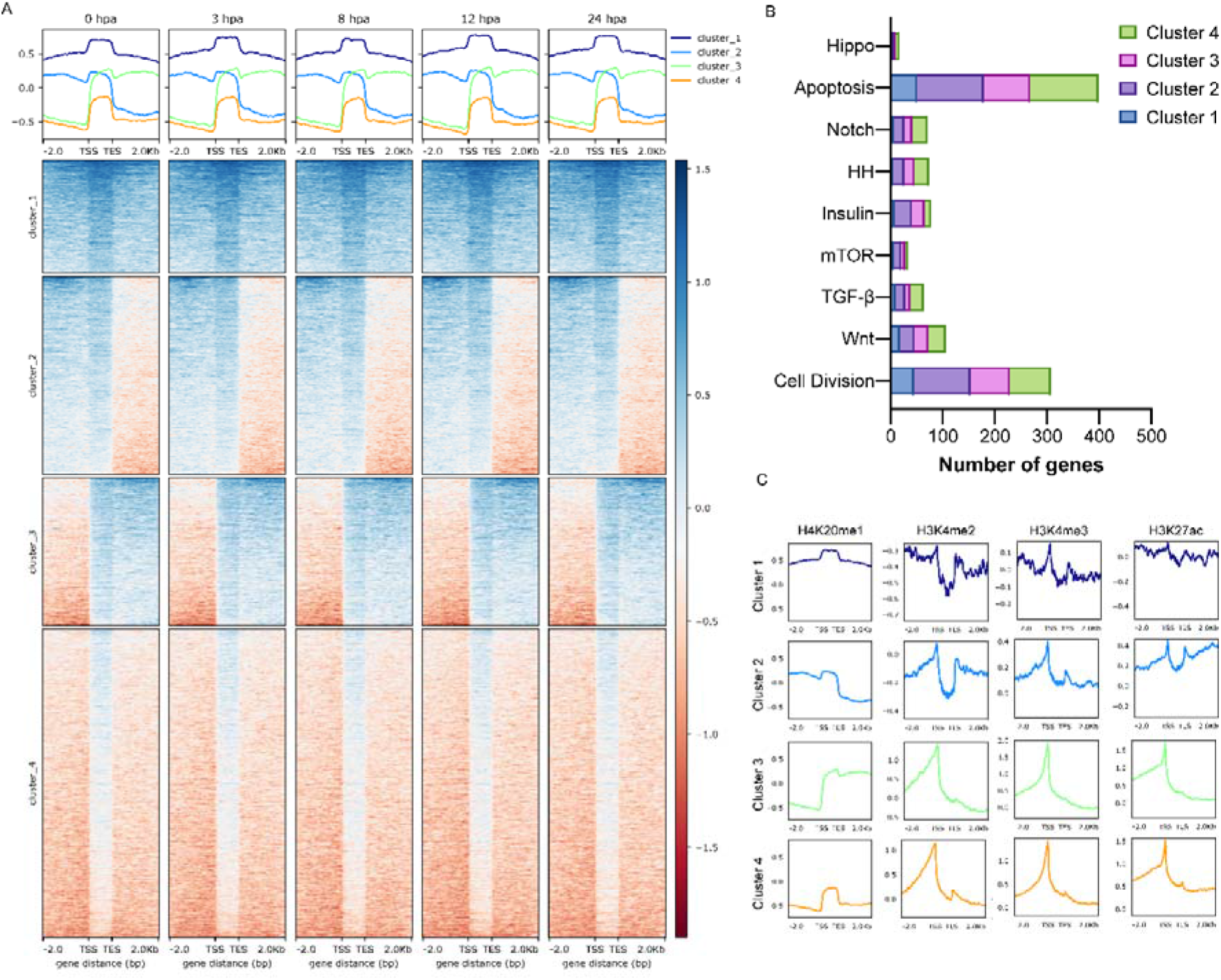
H4K20me1 occupancy during head regeneration indicates a negative association with activation associated histone modifications, open chromatin, and active transcription. A) The occupancy of H4K20me1 on gene bodies across different regenerating time points following head amputation. B) Representation of pathways in the four clusters obtained based on the occupancy of H4K20me1 during regeneration. C) The four clusters obtained based on the occupancy of H4K20me1 on gene bodies were used to investigate the occupancy of the three activation-associated histone marks: H3K4me2, H3K4me3, and H3K27ac at the 0 hpa timepoint. D–F) The four clusters have been used to interrogate the chromatin accessibility and the transcription of the respective genes. The ATAC-seq reads and the RNA-seq reads during a regeneration time course have been plotted for the four clusters separately and represent the openness of the genomic regions and the resultant transcription from those regions, respectively.

The readout of active transcription is the resulting gene expression, which is mediated by opening compacted chromatin. The available ATAC seq (Assay for Transposase-Accessible Chromatin using sequencing) data allows us to investigate the chromatin accessibility during regeneration (Cazet et al. 2021) and the RNA sequencing data is the final readout for gene regulation. The cluster 1 genes have the highest level of H4K20me1, the least intensity of the ATAC seq reads, and the lowest levels of transcription. Cluster 2 genes that have a low level of H4K20me1 at the promoter show slightly higher accessibility and similarly correlated transcription. Clusters 3 and 4, which have clear depletion of H4K20me1 at the promoter regions, show the highest level of openness in the chromatin and the highest level of transcription compared to all the four clusters (Fig. 6).

**Fig. 6:**
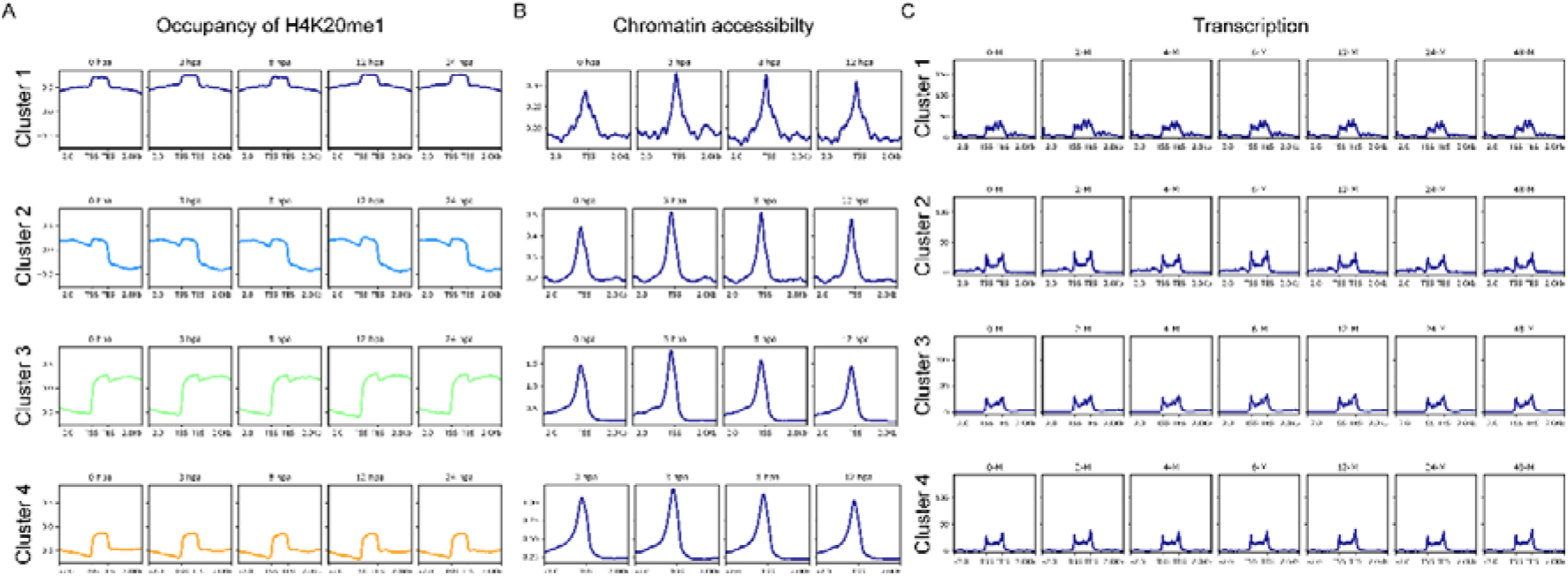
H4K20me1 excludes transcriptionally active regions of the *Hydra* genome. The four clusters obtained based on the occupancy of H4K20me1 on gene bodies have been used to understand the chromatin accessibility and the transcription of the respective genes. The ATAC seq reads and the RNA seq reads during a regeneration time course have been plotted for the four clusters separately and represent the openness of the genomic regions and the resultant transcription from those regions, respectively.

## DISCUSSION

Regeneration recapitulates developmental patterning and is aided by extensive changes in chromatin architecture and consequently gene expression. These changes are regulated by various epigenetic modifiers and we sought to identify the methylome-modulators in *Hydra* axis patterning and regeneration. Upon chemical inhibition, we identified a significant role for SETD8 in both head and foot regeneration. Inhibition of SETD8 led to a transient reduction of the target H4K20me1 histone modification which results in delayed head regeneration. The emergence of tentacles is delayed by 12 hours, and the polyps are unable to complete differentiation of all the tentacles within 72 hours. Foot differentiation is also severely impacted following the chemical treatment. In *Hydra*, the Wnt signalling pathway and activity of β-catenin are required in the early stages of regeneration of both the head and the foot (Gufler et al. 2018). We investigated the process of head regeneration in deeper detail to understand the mechanism of SETD8 function in *Hydra* physiology. The earliest step of regeneration, wound healing, is morphologically not affected by the inhibition of SETD8. However, at a molecular level, various Wnt signalling components, apoptosis-related genes, Jun-kinase, and many insulin signalling components are dysregulated upon inhibition of SETD8. Brachyury is a bonafide head specification marker among the various genes that are part of the Wnt signalling pathway and direct targets of the TCF7L2 transcription factor. The formation of the head organizer by expression of various patterning transcription factors like *Brachyury* is dramatically altered when SETD8 is inhibited. Impairment of both the head and foot regeneration by inhibiting the SETD8 enzyme indicates a role in the position-independent function of the Wnt signalling pathway. SETD8 in *Hydra* is upregulated at both mRNA and protein levels upon activating the Wnt signalling pathway, and physically interacts with β-catenin in the nucleus. Therefore, similar to other systems where the SETD8 methyltransferase has been studied in the context of organismal development (Li et al. 2009; Huang et al. 2021), in *Hydra*, where the axis patterning processes are continuously active, SETD8 interacts with the β-catenin in the nucleus and possibly has a role in transcriptional regulation.

To decipher the mechanisms underlying transcriptional regulation by SETD8, we characterized the target histone modification, H4K20me1. This modification is present in *Hydra* and can be detected by available antibodies against the mammalian histone mark allowing us to study its dynamics. H4K20me1 is present all across the body column of *Hydra* but has a slightly higher localization near the hypostome and the foot of the polyp. These are two regions of the polyp with greater numbers of differentiating cells and the most patterning events taking place. The higher presence of the histone mark indicates a putative role in these processes. When the body axis is perturbed by ectopic activation of the Wnt signalling pathway, the whole polyp acquires head-like characteristics, and tentacles arise all over the body column. In these conditions, we identified an enhanced occupancy of H4K20me1 at the promoter regions of a homeodomain transcription factor Margin, an established direct target of β-catenin (Reddy et al. 2019). In addition, the regulation of SETD8 expression is also associated with the presence of H4K20me1 at its promoter, indicating has a self-regulatory role for SETD8 downstream of the Wnt signalling pathway. We have identified the molecular mechanism of action of SETD8 where, upon switching on the Wnt signalling pathway, β-catenin translocates into the nucleus. This leads to the interaction of SETD8 and β-catenin at the TCF binding regions of the genome and culminates in transcriptional activation, as seen for both the validated targets *Margin* and *setd8* (Fig. 7).

**Fig. 7:**
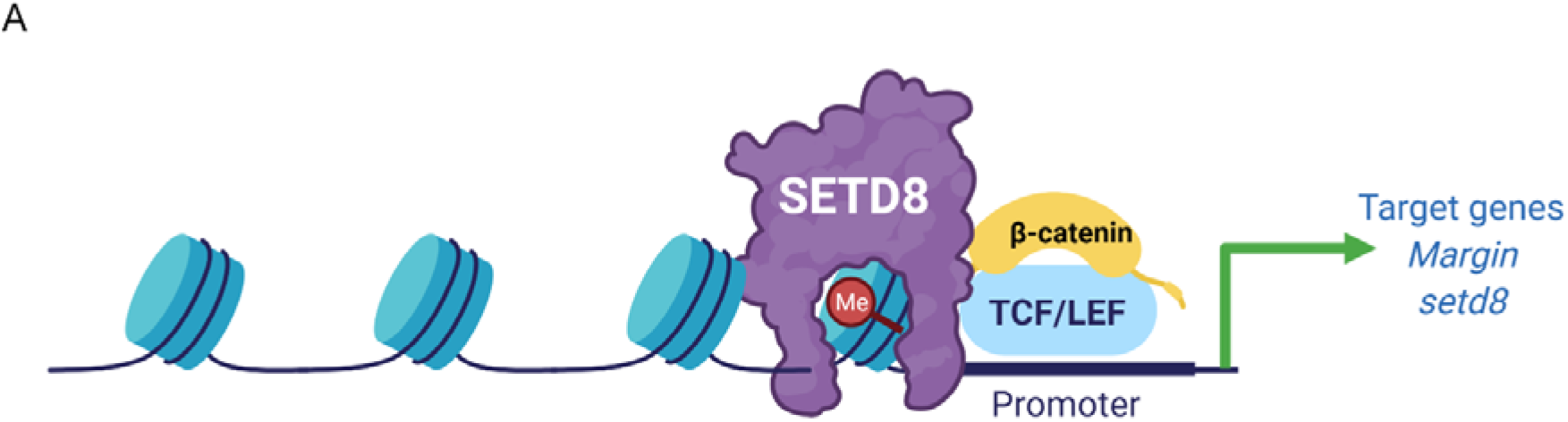
Molecular mechanism of action of SETD8 in *Hydra*. Upon activation of the Wnt signaling pathway, β-catenin is translocated to the nucleus, where it interacts with the TCF7L2 transcription factor to activate the transcription of target genes. The effective activation of the signaling pathway is correlated with SETD8 interacting with β-catenin at the promoters of the target genes and depositing the H4K20me1 mark on the nucleosomes

Investigation of the global occupancy of H4K20me1 upon ectopic activation of the Wnt signalling pathway revealed that H4K20me1 excludes sites occupied by the transcription activation associated histone marks described in a previous study (Reddy et al. 2020). When the axis of the organism is perturbed, there is a modest increase in the occupancy of H4K20me1 globally while during regeneration, the H4K20me1 occupancy does not drastically change across time points. The occupancy is observed on many genes associated with the different cellular and physiological processes and differentially represented in various signalling pathways.The readout of active transcription is the resulting gene expression, and the opening of the chromatin mediates this. The available ATAC-seq data allows us to investigate the chromatin accessibility during regeneration and the RNA sequencing data is the final readout for gene regulation. In addition to the histone modifications, the occupancy of H4K20me1 is also excluded from open chromatin regions displaying active transcription. The occurrence of H4K20me1 extends downstream of the promoters and into gene bodies, indicating an association with transcriptional elongation. H4K20me1 occupancy is known to be associated with the elongation of rapidly transcribing genes in mammalian cells (Vakoc et al. 2006; Veloso et al. 2014). Contrastingly, in senescent cells and neurons, H4K20me1 occupancy seems to prevent elongation (Wang et al. 2015; Tanaka et al. 2017). Thus, the effect of H4K20me1 occupancy on transcription appears to be highly context-specific and regulated in a cell type-specific and location-specific manner.

The role of SETD8 and H4K20me1 in transcriptional regulation is not well established. At promoters of genes involved in erythropoiesis, loss of SETD8 causes reduced chromatin accessibility and impaired differentiation (Myers et al. 2020). In zebrafish and *Drosophila*, H4K20me1 is positively correlated with transcription in a few developmental contexts involving the Wnt signalling pathway (Li et al. 2011; Huang et al. 2021). This has also been observed in global high-resolution profiling done in human T cells (Barski et al. 2007). We observe this positive correlation with transcriptional regulation in the Wnt/β-catenin signalling network in *Hydra*, and the delay in regeneration can be explained by a reduction specifically in the Wnt-mediated differentiation potential of the *Hydra* cells to regenerate lost head and foot structures. A contrasting role for this modification has also been identified using *in vitro* studies involving the L3MBT1 protein, which is critical to transcriptional repression (Kalakonda et al. 2008). During the progression of cell cycle stages, the SETD8 protein is necessary for maintaining chromatin condensation and acts via the PCNA axis (Abbas et al. 2010). This is important to maintain the repressed state of transcription to allow mitosis to move forward and is additionally facilitated by the reduction in the levels of the demethylase PHF8 in the prophase of cell division (Karachentsev et al. 2005; Liu et al. 2010). However, in the context of *Hydra* head regeneration, where there is a necessity for various signalling pathways to interact, the trend indicates that H4K20me1 is negatively correlated with the transcription of genes. The regeneration process involves various cellular processes, one of which is the reduced cellular proliferation and increased transdifferentiation in the morphallactic regeneration of *Hydra*. Therefore, the H4K20me1 mark at a global scale may show an association with reduced transcription, similar to its role in cell cycle regulation. While SETD8 is conserved in flies and humans, the enzyme and the target H4K20me1 have distinct roles in the eye development of *Drosophila* (Crain et al. 2022). The transient decrease in the target mark in *Hydra* and the marked phenotype observed also point towards such a role in the Cnidarian.

Therefore, this mark’s dual mode of action is more specific than we previously could decipher. The specificity is at TF binding sites, and a deeper investigation of the TCF binding sites displayed distinct regulation of H4K20me1 at these motifs in response to Wnt signalling activation. We have, therefore, identified a putative repressive mark in *Hydra* downstream of the H4K20me1 mark. The typically pericentric H4K20me2 and me3 marks may have a transcriptional regulatory role in this early eumetazoan which evolved to be restricted only to pericentromeric heterochromatinization in higher animals. The unique physiology of *Hydra* enables studying and visualising the different modes of action for the histone post-translational modification, H4K20me1, and understand its dual nature in transcriptional regulation (Fig. 8).

**Fig. 8:**
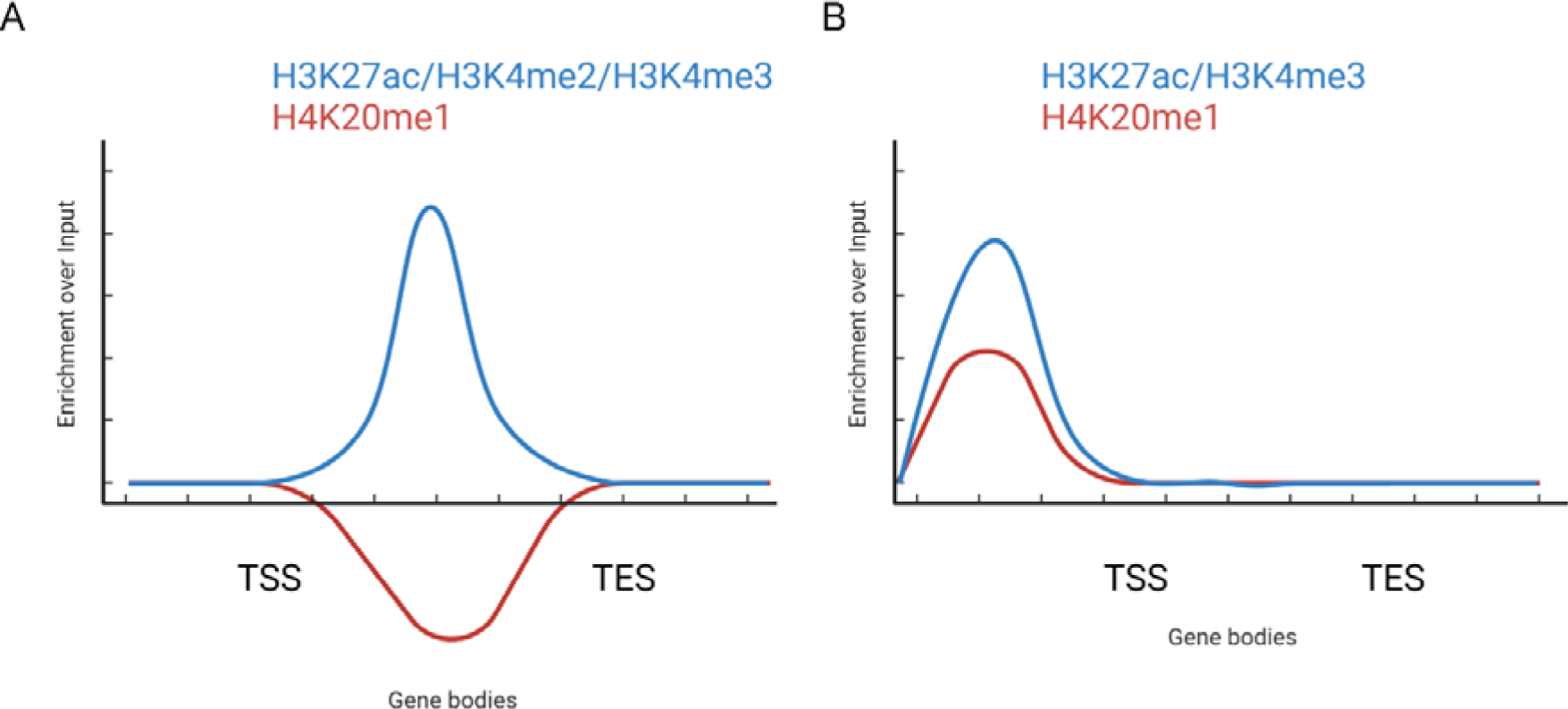
The histone mark H4K20me1 plays a dual role in *Hydra*. A) Based on our global analysis of gene bodies, during regeneration and ectopic activation of Wnt signaling, the occupancy of H4K20me1 is negatively correlated with the occupancy of the activation associated histone marks. B) Contrastingly, based on our localized analysis on the promoters of highly upregulated genes in the context of Wnt signaling mediated axis patterning, H4K20me1 acts as an activator of gene expression and its occupancy coincides with that of the activation associated histone marks.

## MATERIALS AND METHODS

### *Hydra* culture

A clonal culture of *Hydra vulgaris* Ind-Pune was maintained at 18°C in Hydra medium (HM) (100 µM KCl, 100 µM MgSO_4_.7H_2_O, 1 mM CaCl_2_.2H_2_O, 1 mM Tris (pH-8.0), 1 mM NaCl) using standard protocols described previously (Horibata et al. 2004). *Hydra* polyps were fed with freshly hatched *Artemia nauplii* larvae daily and cleaned 6–8 h post-feeding.

### Inhibitor treatments

The details of the inhibitors used and the treatment regime as shown in Fig. 1B are provided in the Supplementary Information.

### Regeneration assays

*Hydra* polyps were treated with target inhibitors for 8 h and amputated. Following amputation, the inhibitor treatment was continued till the target time points and assayed for head and foot regeneration. The detailed methods are given in the Supplementary Information.

### Whole-mount in situ hybridization on regenerating tips

*Hydra* polyps were treated with target inhibitors for 8 h and decapitated. Following decapitation, the inhibitor treatment was continued till the target time points of 1 hour post amputation (hpa), 2 hpa, 4 hpa, and 8 hpa. The polyps were then processed for whole-mount *in situ* hybridization as described previously (Reddy et al., 2020), and the detailed method is given in the Supplementary Information.

### RNA sequencing

The polyps were decapitated, and the regenerating tips were collected at 0 hpa, 3 hpa, 8 hpa, 12 hpa, and 24 hpa. RNA was isolated using Trizol and used to perform 3’ mRNA sequencing using the QuantSeq 3’ mRNA-Seq Library Prep Kit FWD for Illumina (Lexogen, Austria) according to the manufacturer’s instructions.

### Chromatin immunoprecipitation (ChIP)

Two thousand *Hydra* polyps per time point were decapitated, and the regenerating tips were collected at 0 hpa, 3 hpa, 8 hpa, 12 hpa, and 24 hpa to perform ChIP. Two thousand *Hydra* polyps per treatment condition were used for ALP treatment and fixed for ChIP. ChIP was performed as described previously (Reddy et al. 2020) and the detailed method is given in the Supplementary Information.

### ChIP-Seq library preparation and sequencing

A total of 1 ng of purified ChIP-ed DNA for each sample was used for library preparation using Ultra II DNA kit (NEB) as per the manufacturer’s instructions. Library concentration was determined using Qubit (Thermo Scientific, USA), and average fragment size was estimated using DNA HS assay on Bioanalyzer 2100 (Agilent Technologies, USA) before pooling libraries at equimolar ratio. 1.5 pM of the denatured libraries were used as an input to obtain sequencing reads using Nextseq 550 (Illumina) at IISER Pune. The detailed protocol is provided in the Supplementary Information.

### ChIP-Seq Analysis

The quality of the fastq files was checked using FastQC (Andrews 2010), and the quality control (QC) data was consolidated using MultiQC (Ewels et al. 2016). The fastq files were aligned to the latest assembly of the *Hydra* genome using the bowtie2 aligner (Langmead and Salzberg 2012). The sam files were converted to bam files and indexed using samtools (Li et al. 2009). The sorted bam files were used to generate bigwig files using deeptools (Ramirez et al. 2016). The bam files for the H4K20me1 ChIP samples were normalized against the respective input bam file for each timepoint. The combinatorial occupancy matrices were generated using the computeMatrix tool and the plots were generated using the plotHeatmap tool in the deeptools package. The detailed code is provided in the Supplementary Information.

Further detailed methods are provided in the Supplementary Information, and the raw data will be provided upon request to the reviewers.

## DATA AVAILABILITY

The data discussed in this publication have been deposited in NCBI’s Gene Expression Omnibus (Edgar et al. 2002; Barrett et al. 2013) and are accessible through GEO Series accession number GSE205918 (https://www.ncbi.nlm.nih.gov/geo/query/acc.cgi?acc=GSE205918).

## ACKNOWLEDGEMENTS

We thank all Galande Lab members and especially Ankita Sharma for the discussions. We thank Ankita Sharma for help with the sequencing runs. We thank Amarendranath Soory for the discussions and feedback. The work was supported by internal grants from Indian Institute of Science Education and Research, Pune and Shiv Nadar University (Delhi, NCR) to SG. AG was supported by fellowship from the CSIR. SS is supported by fellowship from the CSIR.

## Author contributions

AG and SG conceived the project and wrote the manuscript. AG, MP, and SS performed experiments. AG performed the ChIP-Seq, ATAC-seq and transcriptome analyses. AG and SG interpreted the data. SG supervised the project. All authors read and approved the final manuscript.

