## Supplementary_Information for "H4K20me1 plays a dual role in transcriptional regulation during regeneration and axis patterning of *Hydra*" (from another version of this preprint: version 1, 2022-06-01)

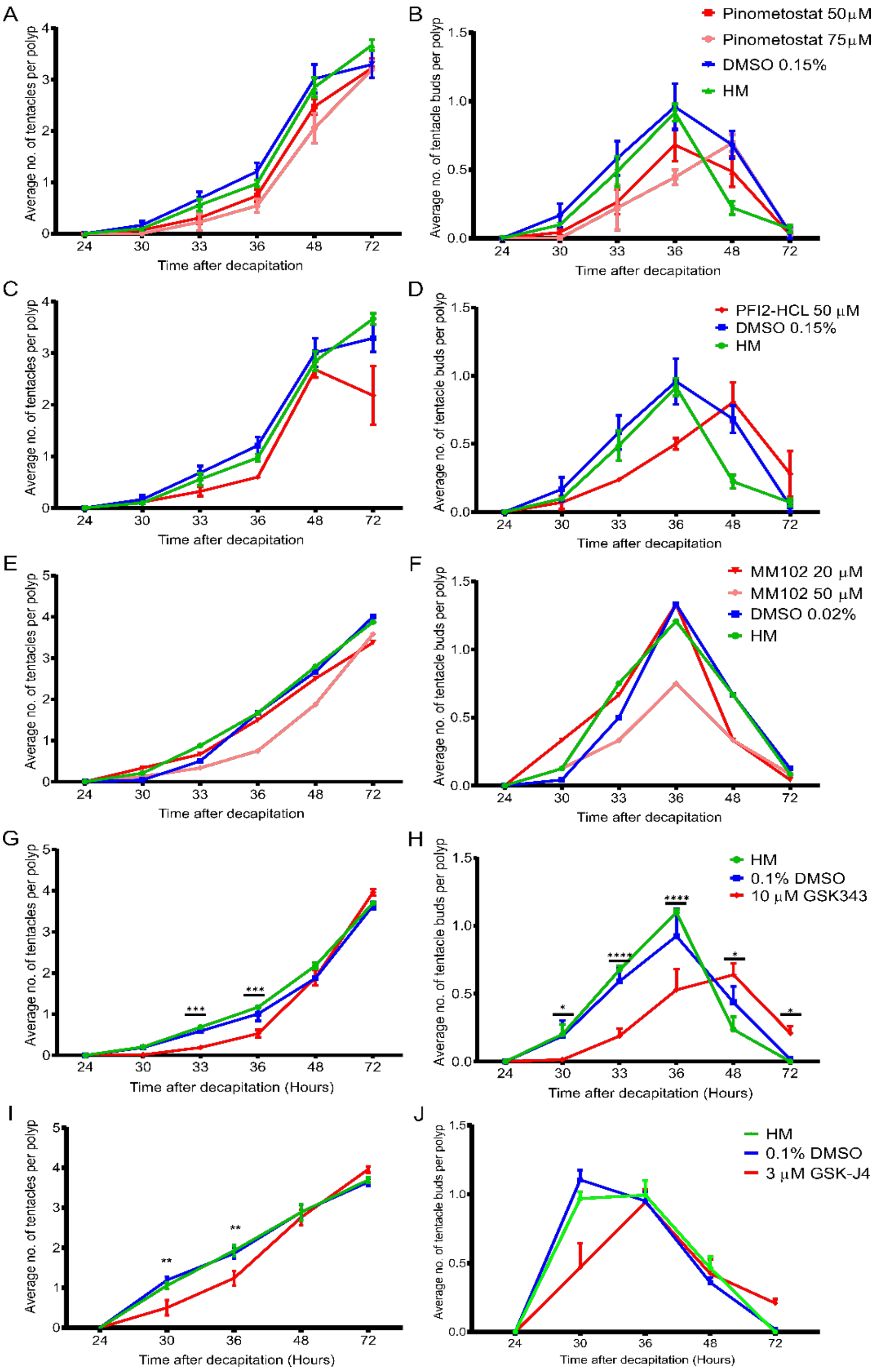


**Supplementary Figure S1:** Screening with various chemical inhibitors against epigenetic molecules to determine the effect on regeneration. The regeneration assays were conducted by counting the number of tentacles and tentacle buds in each polyp after amputation, across multiple experiments with various inhibitors. The graph depicts the average number of tentacles (left) and the number of tentacle buds per polyp (right) at each time point post decapitation during the course of regeneration. (N = 5, n = 25, *- P <0.5, ***- P < 0.001, ****- P <0.0001).

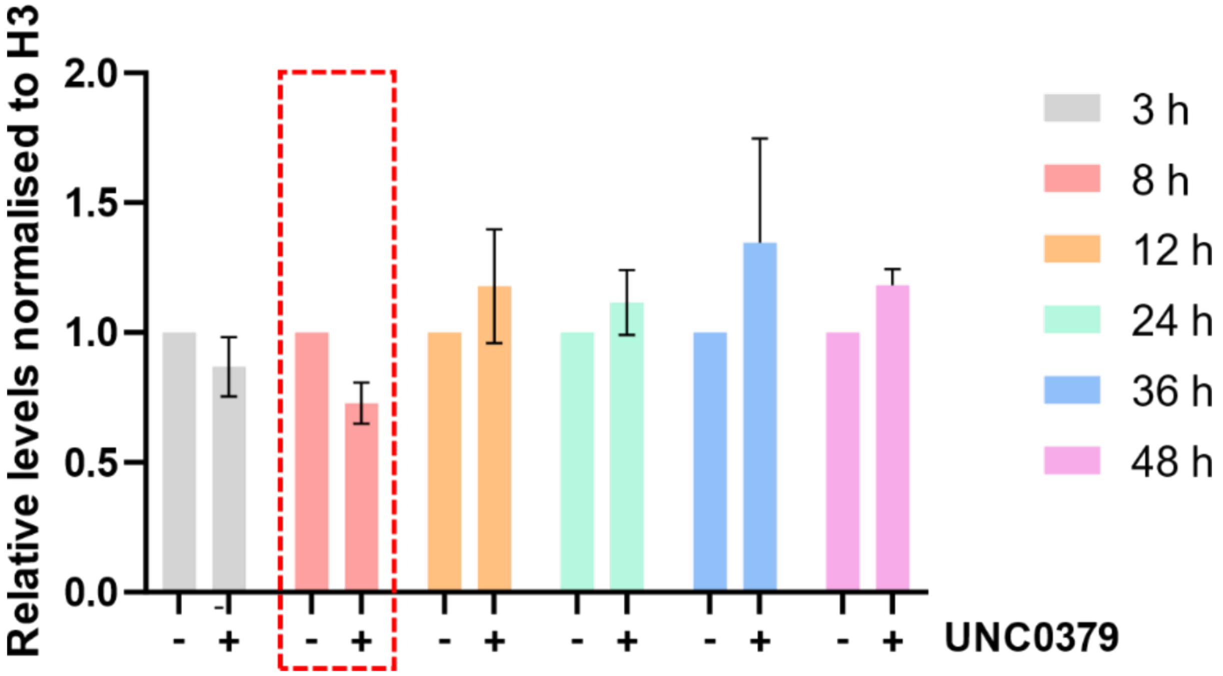


**Supplementary Figure S2 :** Effect of SETD8 inhibitor, UNC0379, on H4K20me1. Quantitation of bands resulting from a Western blot against H4K20me1 normalized to total histone H3 in the polyps. The various time points of chemical treatment are indicated in the legend. There is a marked reduction of H4K20me1 after 8 h of chemical treatment highlighted by the red dashed box. Histone H3 was used as a normalizer for detecting relative enrichment of H4K20me1.

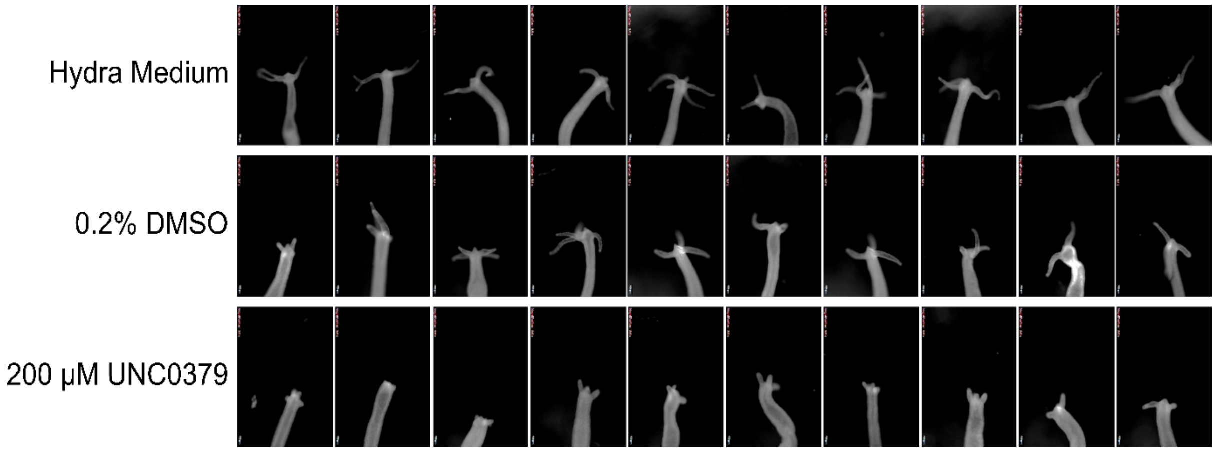


**Supplementary Figure S3** : Regeneration rate and morphology of tentacles upon SETD8 inhibitor treatment. The polyp morphology shows that polyps growing in *Hydra* medium and 0.2% DMSO show no difference in tentacle development, however, inhibition of SETD8 significantly delays the regeneration of tentacles. The stunted tentacle buds in the lower panel indicate the effect of UNC0379.

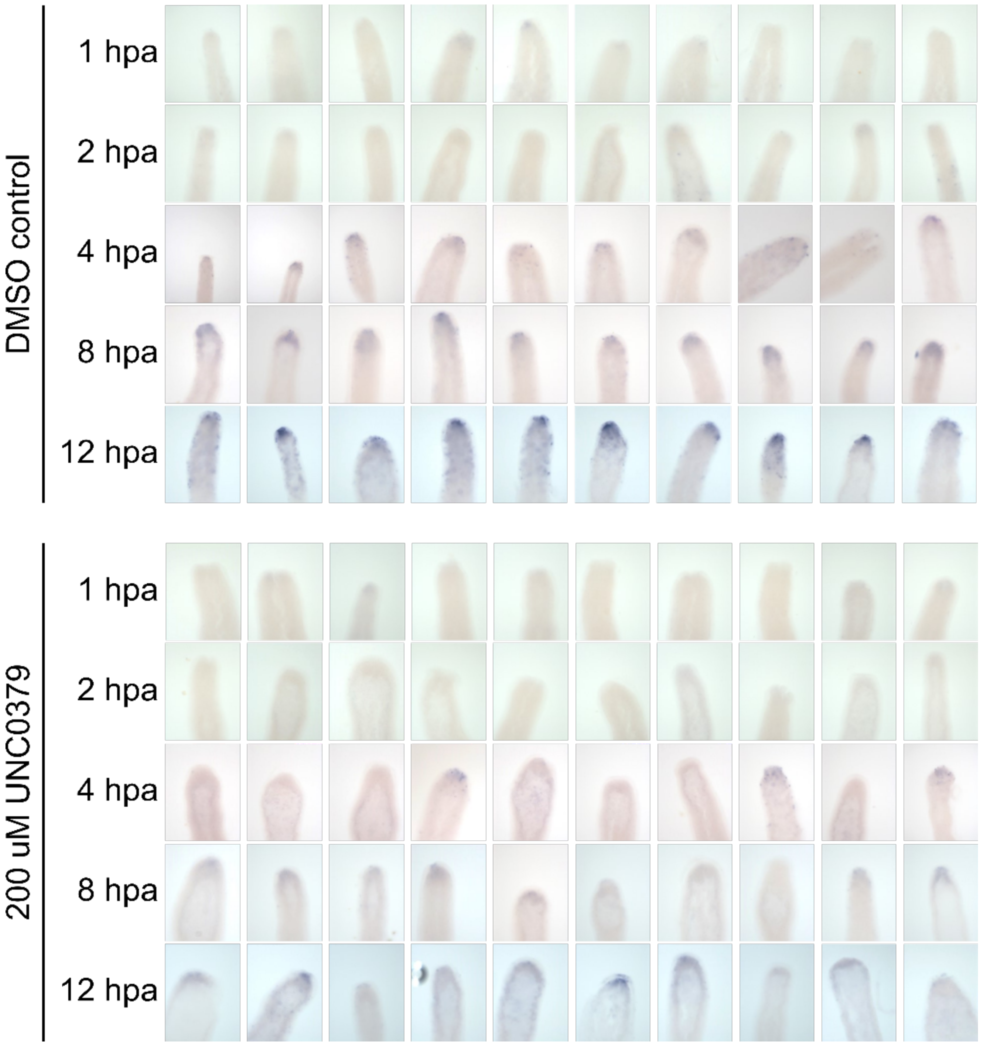


**Supplementary Figure S4 :** Whole mount RNA in situ hybridization against the *Brachyury1* gene in regenerating polyps at 1, 2, 4, 8 and 12 hours post amputation (hpa), with and without inhibitor treatment. The blue stain depicts the expression pattern of *Brachyury1* and shows that the inhibitor treatment impairs the expression of *Brachyury1*. Scale bar measures 100 µm.

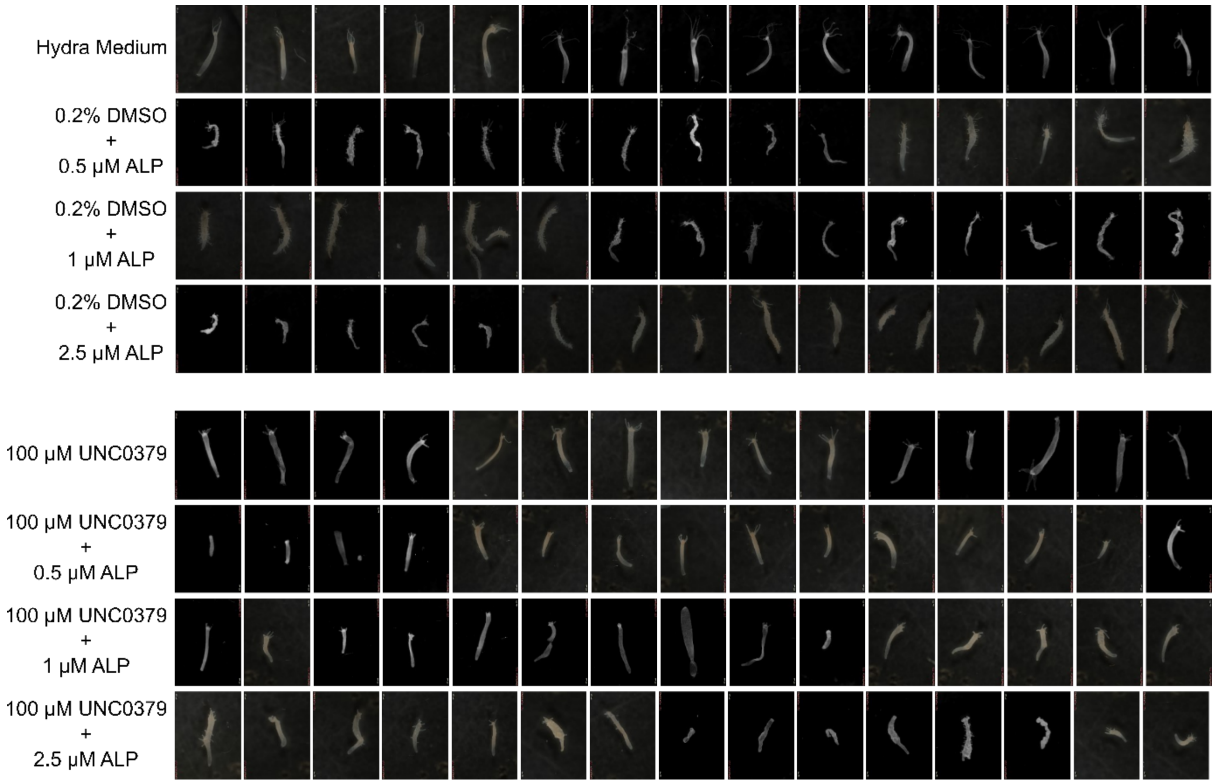


**Supplementary Figure S5:** Combinatorial treatment of polyps with Alsterpaullone and UNC0379 was performed. Polyps were treated in an increasing concentration of ALP (0.5 $\mu$M, 1 $\mu$M and 2.5 $\mu$M) and displayed increasing severity of the multi-tentacle phenotype with ALP concentration. DMSO as a vehicle was used at 0.2% concentration and the SETD8 inhibitor UNC0379 at 100 $\mu$M concentration. The ectopic tentacle phenotype was rescued upon treatment with the SETD8 inhibitor.

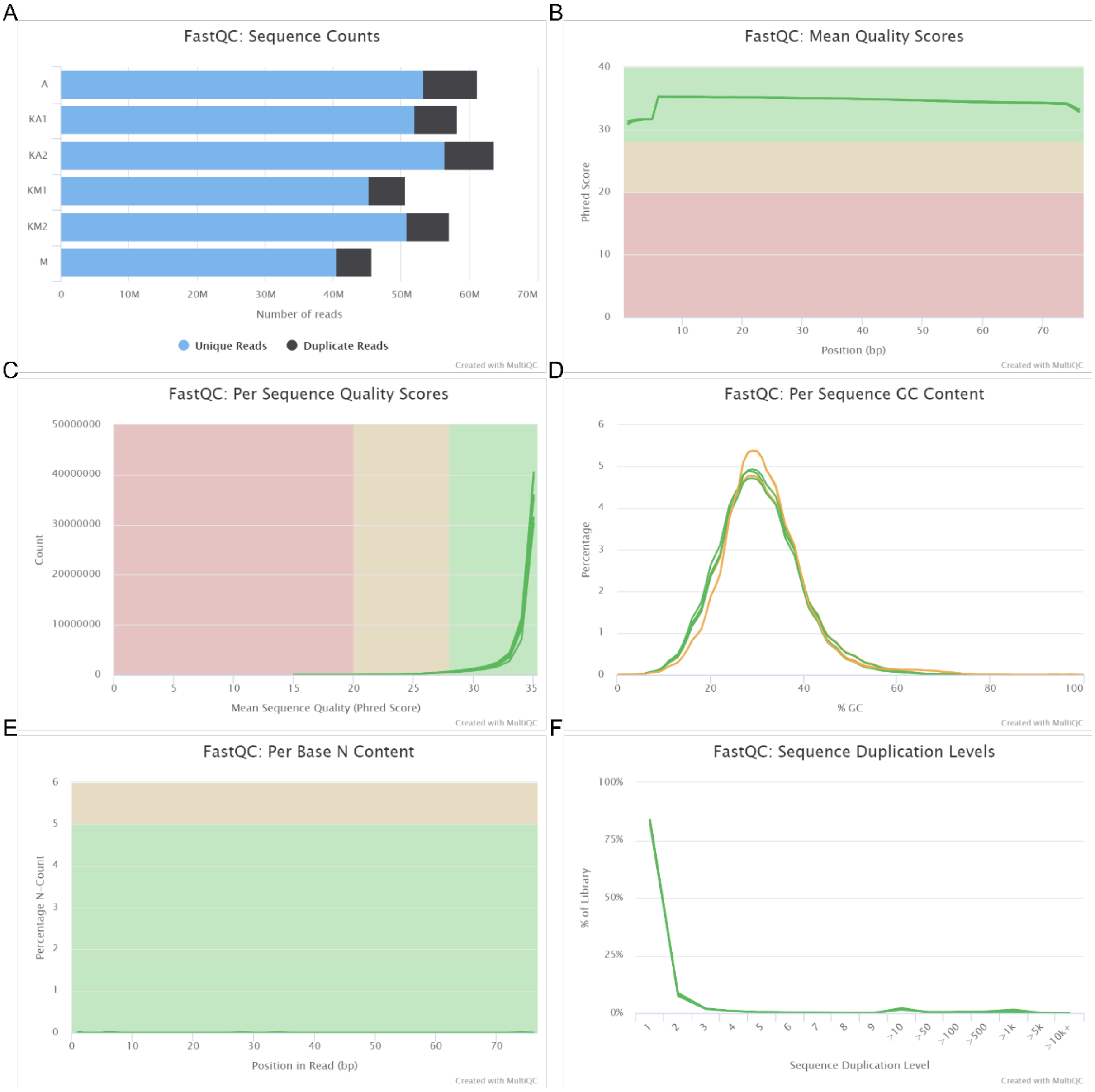


**Supplementary Figure S6**: Quality profile of sequencing reads from H4K20me1 ChIP-seq using ALP treated polyps generated using FastQC tool. M indicates Master control and A stands for ALP treated polyps. (A) Sequence counts for each sample. Duplicate read counts are an estimate only. (B) The mean quality value across each base position in the read indicated by Phred scores. (C) The number of reads with average quality scores. (D) The average GC content of reads. The GC content of the sequencing reads reflects the genomic GC content of *Hydra vulgaris* (~30%). (E) The percentage of base calls at each position for which an 'N' was called. There is a negligible percentage of 'N' bases called indicating a good efficiency of base calling in the sequencing run. (F) The relative level of duplication found for every sequence. The plots show very less over-sequencing of any one sample.

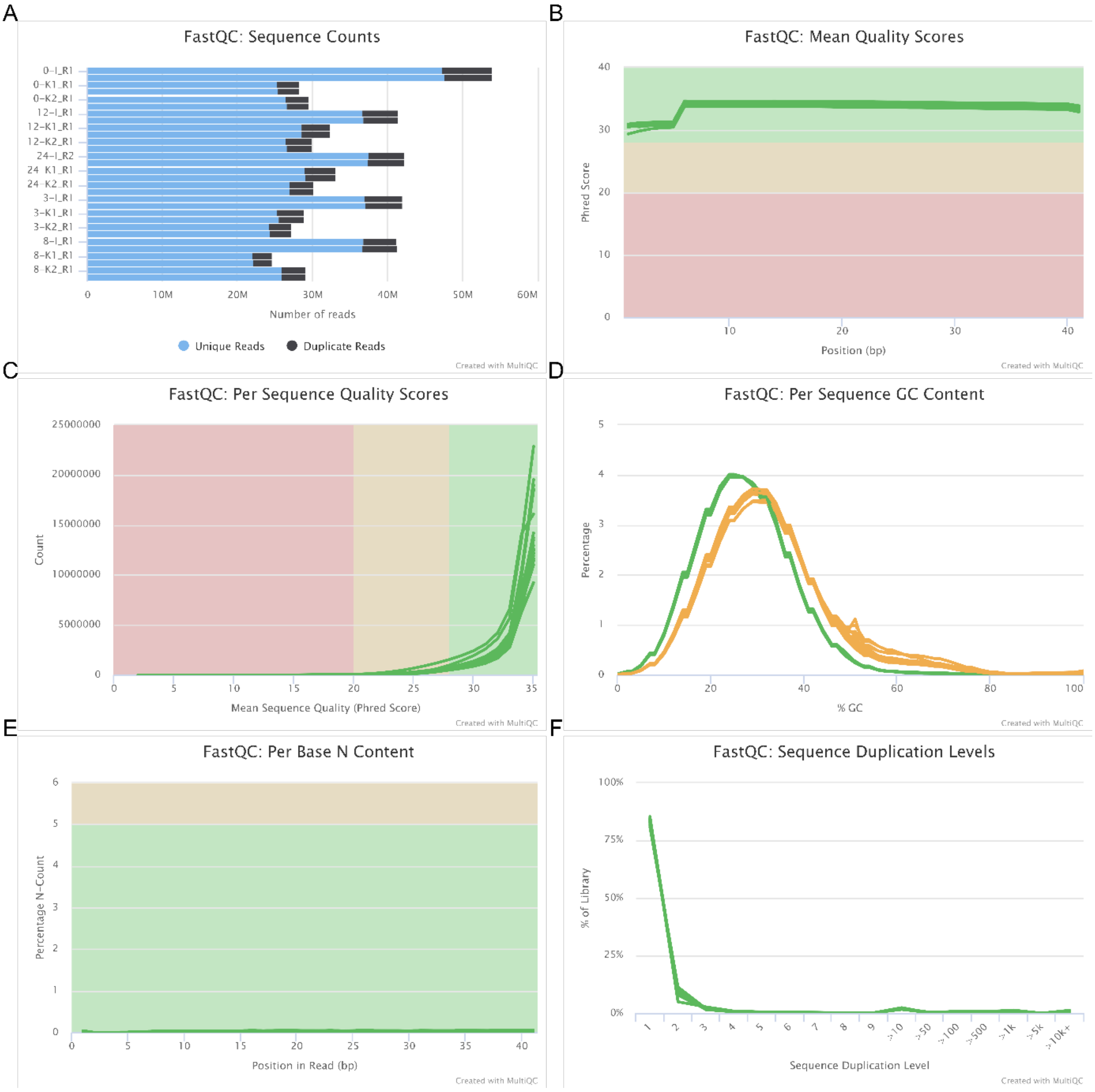


**Supplementary Figure S7:** Quality profile of sequencing reads from H4K20me1 ChIP-seq using regenerating tips of polyps generated using FastQC tool. (A) Sequence counts for each sample. Duplicate read counts are an estimate only. In all our samples, we obtained a good percentage of unique reads with the average duplicate reads being less than 12% across all the samples. (B) The mean quality value across each base position in the read. (C) The number of reads with average quality scores. (D) The average GC content of reads. The GC content of the sequencing reads reflects the genomic GC content of *Hydra vulgaris* (~30%). (E) The percentage of base calls at each position for which an 'N' was called. There is a negligible percentage of 'N' bases called indicating a good efficiency of base calling in the sequencing run. (F) The relative level of duplication found for every sequence. The plots show very less over-sequencing of any one sample.

### Supplementary Methods

#### Inhibitor treatments

To perform the inhibitor treatments, the following inhibitors were used: Pinometostat (EPZ5676) for the KMT4 enzyme (Cat # S7062, Selleckchem, Houston, Texas), MM-102 for the KMT2A enzyme (Cat # S7265, Selleckchem, Houston, Texas), UNC0379 for the KMT5A enzyme (Cat # S7570, Selleckchem, Houston, Texas; Cat # 16400, Cayman Chemicals Co., Ann Arbor, Michigan), PFI-2 HCl for the KMT7 enzyme (Cat # S7294, Selleckchem, Houston, Texas), GSK-J4 for the enzyme KDM6A/6B (Cat # SML0701-5MG, Sigma-Aldrich, St. Louis, Missouri), GSK-LSD1 for the enzyme KDM1A (Cat # SML1072-5MG, Sigma-Aldrich, St. Louis, Missouri), and GSK343 for the enzyme KMT6A (Cat # SML0766, Sigma-Aldrich, St. Louis, Missouri). To identify the working concentration of each inhibitor, 50 *Hydra* polyps were treated for 8 hours (h) (overnight), and the concentration at which 50% of the polyps were disintegrated was identified ($LD_{50}$). A concentration below the $LD_{50}$ was used to perform the regeneration assays.

#### Head regeneration assay

*Hydra* polyps were treated with target inhibitors for 8 h and decapitated. Following decapitation, the inhibitor treatment was continued till 72 h with fresh addition of inhibitor-containing Hydra medium every 24 h. At the time points — 24 h, 30 h, 33 h, 36 h, 48 h, and 72 h — the number of tentacle buds and tentacles were manually counted and recorded. The statistical analysis was performed using the GraphPad Prism version 7.0c for MacOS (GraphPad Software, San Diego, California USA, www.graphpad.com).

#### Foot staining assay

*Hydra* polyps were treated with target inhibitors for 8 h, and the foot was amputated. Following amputation, the inhibitor treatment was continued till 36 h with fresh addition of inhibitor-containing Hydra medium every 24 h. At the target timepoints of 22 h, 26 h, 30 h, and 36 h post-amputation, the polyps were relaxed with 2% urethane/HM for 2 min and fixed using 4 % PFA/PBS for 1 h at room temperature. After fixation, specimens were given three 10 min washes in 1X PBS + 0.25% Triton X-100 (PBST). The animals were then incubated for 15 min at room temperature in 1X PBST containing 0.02% diaminobenzidine (DAB) and 0.003% hydrogen peroxide ($H_2O_2$). After incubation, they were rinsed for 30 min in PBST, mounted on slides in phosphate-buffered glycerol, and examined with a Zeiss microscope.

#### Whole-mount in situ hybridization on regenerating tips

*Hydra* polyps were treated with target inhibitors for 8 h and decapitated. Following decapitation, the inhibitor treatment was continued till the target time points of 1 hour post amputation (hpa), 2 hpa, 4 hpa, and 8 hpa. The polyps were then relaxed by treatment with 2% urethane/HM for 2 min and fixed using 4 % PFA/HM at 4°C overnight. Digoxigenin labelled RNA probes for *Hv_Bra1* were prepared by in vitro transcriptions from templates amplified from a recombinant pCR Blunt II TOPO (Cat # 450031, Invitrogen™, Waltham, Massachusetts) plasmid containing the *Hv_Bra1* gene using PCR. (DIG Labelling Mix, Cat #

1277073910, Sigma-Aldrich, St. Louis, Missouri; SP6 RNA Polymerase, Cat # 10810274001, Sigma-Aldrich, St. Louis, Missouri; T7 RNA Polymerase, Cat # 10881767001, Sigma-Aldrich, St. Louis, Missouri). Whole-mount in situ hybridization was performed on the polyps as described previously (Martinez et al., 1997) with the following changes. Treatment with proteinase-K was performed for 5 min, and heat-inactivation of the endogenous alkaline phosphatases was done at 70°C for 15 min in 1X SSC. Digoxigenin-labelled RNA probe at a concentration of 150 ng/ml was used for hybridization at 59°C. The post-hybridization washes were performed using 1X SSC-HS gradients. After staining with 50% NTMT/50% BM-purple AP substrate for 1 h at room temperature, the animals were mounted in 80% glycerol for imaging.

**Actin filament staining**

*Hydra* polyps were treated with target inhibitors for 8 h and decapitated. Following decapitation, the inhibitor treatment was continued till the target time points of 0 hours post amputation (hpa), 2 hpa, and 4 hpa. At the target timepoints, the polyps were relaxed with 2% urethane/HM for 2 min and fixed using 4 % PFA/PBS for 1 h at room temperature. After three 10 min washes in 1X PBS and permeabilization in 1X PBS + 0.1% Triton X-100 (1X PBST), staining was done with Alexa Fluor 568 phalloidin (Cat # A12380, Invitrogen™, Waltham, Massachusetts) diluted 1:200 in 1X PBST at room temperature for 1 h. Finally, the polyps were washed in 1X PBST (3 × 10 min), mounted on slides using VECTASHIELD® Antifade Mounting Medium (Cat # H-1000-NB, Novus Biologicals, Littleton, Colorado) and imaged using the ApoTome microscope (Carl Zeiss, Jena, Germany).

**Immunofluorescence assay**

*Hydra* polyps were treated with target inhibitors for 8 h and decapitated. Following decapitation, the inhibitor treatment was continued till the target time points of 0 hours post amputation (hpa), 2 hpa, and 4 hpa. At the target timepoints, the polyps were relaxed with 2% urethane/HM for 2 min and fixed using 4 % PFA/PBS for 1 h at room temperature. Following fixation, the polyps were permeabilized using 1X PBS + 0.1% Triton X-100 (1X PBST) for 30 min with changes every 10 min. Blocking was done with 1X PBST + 20% FBS for 1 h at 4°C. The solution was replaced with a fresh blocking solution containing α-H4K20me1 (rabbit polyclonal IgG, Cat # CS200569, EMD Millipore) primary antibody at a dilution of 1:100 and incubated overnight at 4°C. After washing with 1X PBST thrice for 10 min each, the polyps were incubated in a solution containing secondary antibody at a dilution of 1:100 (anti-rabbit IgG-Alexa568, Cat # A11011, Invitrogen™, Waltham, Massachusetts). After washing, the nuclei were stained with 0.5 μg/ml DAPI (4′,6-diamidino-2-phenylindole) for 10 min at room temperature. Finally, the polyps were washed in 1X PBS thrice for 10 min each, mounted on glass slides using Vectashield mounting medium, and imaged using the ApoTome microscope (Carl Zeiss).

**Co-immunoprecipitation**

To perform co-immunoprecipitation, 100 polyps were collected and washed thrice with Hydra medium. The medium was removed entirely, and the polyps were resuspended in RIPA buffer (50 mM Tris-HCl pH-7.4, 150 mM NaCl, 2 mM EDTA, 0.1 % v/v SDS, 1 % v/v NP-40, 0.5% w/v sodium deoxycholate, 30 mM sodium fluoride, 0.2 mM sodium orthovanadate) and incubated on ice for 30 min. The sample was centrifuged at 4°C and 14000 rpm for 20 min twice, and the supernatant was collected. The amount of protein was quantitated using the Pierce™ BCA Protein Assay Kit. Five micrograms of α-active-β-catenin antibody (Cat # 05-665, Sigma-Aldrich, St. Louis, Missouri) was added to 700 µg of *Hydra* lysate and incubated at 4°C overnight. An equal amount of the appropriate IgG control was also used (Normal mouse IgG, Cat # 12-371, Sigma-Aldrich, St. Louis, Missouri). 5% or 35 µg of protein was used as the input control. Following pull down, the samples were incubated with 20 µl of Dynabeads™ M-280 sheep anti-mouse IgG (Cat # 11202D, Invitrogen™, Waltham, Massachusetts) for 3 h at 4 °C with gentle inverting. The supernatant was discarded, and the beads were washed with RIPA buffer thrice for 10 min each. The beads were then heated in 1X Laemmli buffer at 95°C for 15 mins to denature and retrieve all the bound proteins. This was separated on an SDS-PAGE gel and subjected to Western blot using the α-SETD8 antibody.

**Acid extraction of histones**

Two hundred *Hydra* polyps were homogenized in lysis buffer wash buffer (250 mM sucrose, 50 mM Tris-Cl, 25 mM KCl, 5 mM $MgCl_2$, 0.2 mM PMSF, 50 mM $NaHCO_3$, 0.2% Triton X-100, 45 mM Na-Butyrate, 10 mM β-mercaptoethanol, 1X PIC) and nuclei isolated by centrifugation at 800 g for 15 min at 4°C. Histone proteins were isolated using the acid extraction method and resolved by 18% SDS-PAGE as previously described (Jayani et al., 2010). Briefly, the nuclear pellet was completely resuspended in 600 µl of 1 $NH_2SO_4$ and incubated for 2 h at 4°C. Histones were precipitated overnight at -20°C from the supernatant by adding trichloroacetic acid to a final concentration of 33%. Chilled acetone washes were carried out to avoid carry-over of trichloroacetic acid. Histone pellets were air-dried, resuspended in 1x PBS and resolved by SDS-PAGE.

**Chromatin immunoprecipitation (ChIP)**

The polyps were decapitated, and the regenerating tips were collected at 0 hpa, 3 hpa, 8 hpa, 12 hpa, and 24 hpa to perform ChIP. Two thousand *Hydra* polyps per time point were decapitated and the regenerating tips were collected and cross-linked with 1% methanol free formaldehyde. The fixation reaction was quenched with 150 mM Glycine. The fixed regenerating tips were resuspended in Swelling Buffer (25 mM Tris-HCl pH 7.9, 1.5 mM $MgCl_2$, 10 mM KCl, 0.1% NP-40, 1 mM DTT, 0.5 mM PMSF, 1X Protease inhibitor cocktail/PIC) and homogenized using a Dounce homogenizer (35 strokes) to lyse the cells and release the nuclei. The nuclei were pelleted, resuspended in sonication buffer (50 mM Tris-HCl pH 7.9, 140 mM NaCl, 1 mM EDTA, 1 % Triton X-100, 1% SDS, 0.1% Sodium deoxycholate, 0.5 mM PMSF, 1X PIC), and incubated for 30 min on ice. They were then sonicated to obtain an average chromatin size of 300 bp. Chromatin was pre-cleared using 50 µl of a 50% protein A sepharose (GE healthcare) slurry for 1 h at 4°C with gentle inverting. Immunoprecipitations were carried out in ChIP buffer (16.7 mM Tris-HCl pH 8.0, 167 mM NaCl, 1.2 mM EDTA, 1.1% Triton-X 100, 0.01% SDS, 1X PIC) with Anti-H4K20me1. An appropriate IgG control was also used with inverting at 4°C for 14–16 h. The samples were then incubated with 50 µl of a 50% Protein A sepharose slurry (saturated with 0.5% BSA and 10 mg/ml yeast tRNA) for 3 h at 4°C with gentle inverting. ChIP samples were reverse-crosslinked, and the DNA was purified using Phenol:Chloroform:Isomamyl alcohol based precipitation method. Input chromatin was obtained after pre-clearing, by de-crosslinking and purifying input DNA Phenol:Chloroform:Isoamyl alcohol based precipitation method. Purified DNA was subjected to library preparation for sequencing or used for quantitative PCR. The fold enrichment was calculated using the formula – Fold enrichment = $2^{(Ct\ (Target\ antibody)-Ct\ (IgG))}$.

### ChIP-Seq library preparation

A total of 1 ng of purified ChIPped DNA for each sample was used for library preparation using Ultra II DNA kit (NEB) as per the manufacturer's instructions. All library samples were amplified for 9-10 cycles depending on cycle number estimation by qPCR. Amplified libraries were subjected to double-sided bead purification using one round of 0.5X volume of Hi-prep PCR purification kit (Magbio Genomics, USA) to remove the primer dimers and one round of 1.8X volume to remove the fragments larger than 1000 bp. Library concentration was determined using Qubit (Thermo scientific, USA), and average fragment size was estimated using DNA HS assay on Bioanalyzer 2100 (Agilent Technologies, USA) before pooling libraries at equimolar ratio.

### Sequencing and Adaptor trimming

1.5 pM of the denatured libraries were used as an input to obtain sequencing reads. The DNA was loaded onto a NextSeq 500/550 High-Output v2.5 Kit (75 cycles) (Cat # 20024906, Illumina) and sequenced using 35 cycles for read1, 8 cycles for indexes and 35 cycles for read2 on Nextseq 550 (Illumina) at the sequencing facility of IISER Pune.

Following sequencing, the bcl files obtained were converted to fastq (Illumina®).

```
bcl2fastq -R 210402_NB551653_0077_AHNCFTBGXF -p 10 --output-
dir 210402_NB551653_0077_AHNCFTBGXF/fastq_files --sample-sheet
210402_NB551653_0077_AHNCFTBGXF/SampleSheet.csv
```

The sample-specific files were generated by concatenation of all four sequenced lanes.

```
cat 0-I_S11_L001_R1_001.fastq.gz 0-I_S11_L002_R1_001.fastq.gz
0-I_S11_L003_R1_001.fastq.gz 0-I_S11_L004_R1_001.fastq.gz > 0-
I_R1.fastq.gz;
```

### ChIP-Seq Analysis

The quality of the fastq files was checked using FastQC (Andrews, 2010), and the quality control (QC) data was consolidated using MultiQC (Ewels et al., 2016).

```
fastqc 0-I_R1.fastq.gz;
```

The fastq files were aligned to the latest assembly of the *Hydra* genome using the bowtie2 aligner (Langmead and Salzberg, 2012).

```
bowtie2 -p 10 -x Hydra2-0_bowtie2_index -1 0-I_R1.fastq.gz -2
0-I_R2.fastq.gz -S 0-I_R2.sam;
```

The sam files were converted to bam files using samtools (Li et al., 2009).

```
samtools sort -@ 10 0-I.bam -o 0-I_sorted.bam;
```

The bam files were sorted and indexed.

```
samtools index -b -@ 10 0-I_sorted.bam 0-I_sorted.bai;
```

The sorted bam files were used to generate bigwig files using deeptools (Ramirez et al., 2016). The bam files for the H4K20me1 ChIP samples were normalized against the respective input bam file for each timepoint. The combinatorial occupancy matrices were generated using the computeMatrix tool and the plots were generated using the plotHeatmap tool in the deeptools package.

```
bamCompare -p 10 -b1 24-K2_sorted.bam -b2 24-I_sorted.bam -o
24-K2_overInput.bw -of bigwig --normalizeUsing RPKM --
scaleFactorsMethod None;
```

```
computeMatrix scale-regions -R
/home/ashwin/Akhila/Hydra_transcriptomes_mapping/Hydra2-
0_genemodels.gtf -S 0_overInput.bw 3_overInput.bw
8_overInput.bw 12_overInput.bw 24_overInput.bw -p 10 -o
H4K20me1_all_matrix -b 2000 -a 2000 -bs 10;
```

```
plotHeatmap -m H4K20me1_all_matrix -o Reg_H4K20me1_reg.png --
dpi 600 --kmeans 4 --colorMap RdBu -T
H4K20me1_occupancy_GeneBodies-k4 --samplesLabel "0 hpa" "3
hpa" "8 hpa" "12 hpa" "24 hpa" --outFileNameMatrix
H4K20me1_reg-matrix-k4 --outFileSortedRegions H4K20me1_reg-
k4.bed;
```

```
computeMatrix reference-point -R
/home/ashwin/Akhila/Hydra_transcriptomes_mapping/Hydra2-
0_genemodels.gtf -S 0_overInput.bw 3_overInput.bw
8_overInput.bw 12_overInput.bw 24_overInput.bw -p 10 -o
H4K20me1_all_matrix-TSS -b 2000 -a 2000 -bs 10;
```

```
plotHeatmap -m H4K20me1_all_matrix-TSS -o Reg_H4K20me1_reg-
TSS.png --dpi 600 --kmeans 4 --colorMap RdBu -T
H4K20me1_occupancy_GeneBodies-k4-TSS --samplesLabel "0 hpa" "3
hpa" "8 hpa" "12 hpa" "24 hpa" --outFileNameMatrix
H4K20me1_reg-matrix-k4-TSS --outFileSortedRegions
H4K20me1_reg-k4-TSS.bed;
```
